# Preclinical drug response metric based on cellular response phenotype provides better pharmacogenomic variables with phenotype relevance

**DOI:** 10.1101/2020.12.23.424257

**Authors:** Sanghyun Kim, Sohyun Hwang

## Abstract

**Background and Purpose:** Assessment of drug response is typically performed by constructing a dose-response curve of viability and summarizing it to a representative value. However, this is limited by its dependency on the assay duration and lack of reflections regarding actual cellular response phenotypes. To resolve these limitations, we considered contribution of each response phenotype upon a drug treatment to the overall growth behavior.

**Experimental Approach:** The differential equation of phenotype population dynamics was solved analytically without numerical computation. By using the properly figured population dynamics, we explored how the conventional assessment method itself affects the assessment result of drug response, in the context of drug screening. Alternative phenotype metric was compared with the conventional metrics through evaluation of the publicly available drug response data.

**Key Results:** The conventional assessment showed several limitations in the comparative analysis of drug response: a significant time-dependency, and ambiguities in assessment results based on a dose-response curve. Instead, the alternative phenotype metrics provide time-independent phenotype rates of change, that contain all the information of the drug response at a given dose, and better classification including the mechanism underlying growth inhibition.

**Conclusion and Implications:** The conventional dose-response curve is useful for a visual presentation of overall drug responses upon a certain molecular feature qualitatively. In contrast, the phenotype metric is better for assessing therapeutic effectiveness, and would improve preclinical pharmacogenomic analysis through its relevance to a response phenotype.

**Bullet point summary:** *What is already known:*

- Drug response is investigated by constructing a dose-response curve in wide range.
- Conventional assessment metrics of drug response lack reflections regarding actual cellular response phenotypes.

*What this study adds:*

- Limitations of conventional assessments are due to time-dependency of dose-response curve and customary summarization.
- Phenotype metric evaluate a single dose-response that is time-independent and phenotype-relevant.

*Clinical significance:*

- Phenotype metric would improve pharmacogenomic analysis with better classification and phenotype-relevance of drug response.
- Improvement in preclinical pharmacology would bring better translation and useful information in clinical studies.

## 1 INTRODUCTION

Drug responses in cultured cell lines are essential for identifying molecular features associated with therapeutic effectiveness of the drug, through integration with large genomic data (Wang et al., 2011; Barretina et al., 2012; Garnett et al., 2012; Costello et al., 2014; Iorio et al., 2016; Rees et al., 2016). It has been investigated by constructing a dose-response curve of a certain metric, e.g., viability, and the response curve is summarized into a representative quantity. The common summary factors are the concentration of half maximal inhibition (IC_50_) or half maximal effect (EC_50_) as a measure of potency, the maximal inhibition level (E_max_) as a measure of efficacy, and the area under the curve (AUC) as a measure of overall effectiveness. For better pharmacogenomic analysis, two limitations need to be addressed adequately: the issue of irreproducibility in drug response assays, and the discordance between actual cellular response phenotypes and assessment metrics of drug response. Concerns regarding reproducibility in drug response studies are a practical issue raised repeatedly in preclinical studies (Haibe-Kains et al., 2013; Collins and Tabak, 2014; Freedman et al., 2015). On the other hand, the second limitation is a more general issue; while actual response phenotypes during drug treatment are diverse– including senescence and various forms of cell death–conventional metrics such as IC_50_ do not reflect such responses. Instead, these metrics are simply *de facto* standards for crudely categorizing cell lines as either sensitive or resistant without an underlying theoretical basis.

A recent multi-center study of the NIH Library of Network-Based Cellular Signatures Program (LINCS) on drug response assays (Niepel et al., 2019) well evaluated the irreproducibility issue. A notable point of this study was that a growth rate (Hafner et al., 2016; Harris et al., 2016)– instead of viability–was used as a metric for producing a dose-response curve. Viability always depends on the assay duration, and the duration is a relative factor for the doubling time of each cell line. To correct for this confounder, this study used an alternative metric based on the apparent growth rate, under the assumption of a simple exponential cell growth (Hafner et al., 2016; Harris et al., 2016). Indeed, this metric reportedly improved the reproducibility and performance of pharmacogenomic association studies (Hafner et al., 2017). However, apparent-growth-rate-based metrics still depend on time in numerous cases, as observed in the MCF10A cell line treated with neratinib in the LINCS study (Niepel et al., 2019). This implies that cellular growth behavior is not a simple exponential function.

Here the widespread assumption for cellular growth, a simple exponential function, was reconsidered. We investigated how each cellular response phenotype upon a drug treatment contributes to the overall growth behavior, and solved population dynamics of each phenotype. By using this population dynamics, we explored how the conventional assessment method itself–encompassing the metric for construction of a dose-response curve, the assay duration, and the summary factor–affects the assessment result of drug response and revealed its limitations for comparative analysis of drug response. In contrast, the alternative metric based on phenotype population dynamics produces time-independent characteristic quantities of drug response. This could provide better pharmacogenomic variables relevant to the response phenotype of the cell.

## 2 METHODS

### 2.1 Exploring the Conventional Evaluation of Drug Response

#### Phenotype Parameters and Growth Curve

Growth curves for a certain drug-cell line pair were generated by assigning phenotype rates of changes of the population dynamics model presented in Result section (Figure 1a). The rates of changes (*k_P_* for proliferating cells, *s* for senescent cells, and *k_d_* for dead cells) were assumed respectively as Hill function of dose, that is the most widely used functional form for dose-response and even for a growth rate(Hafner et al., 2016).

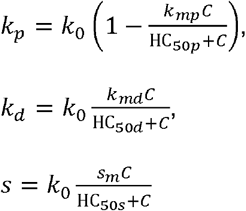

**Figure 1.**
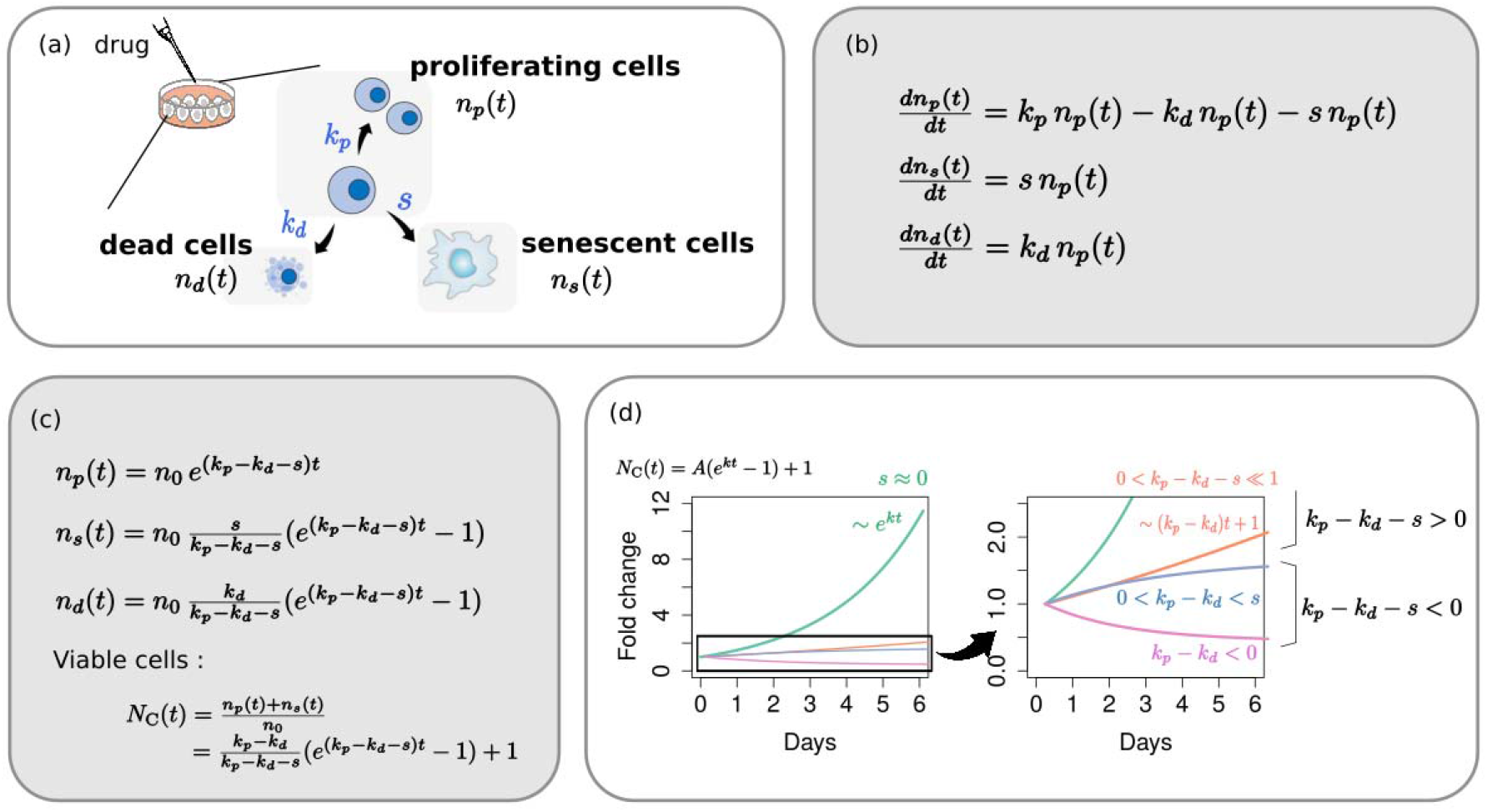
Phenotype dynamics model. (a) A diagrammatic illustration of phenotypic response of the cell upon a drug treatment. Proliferating cells can continue dividing, can enter a state of permanent cell cycle arrest, or can undergo cell death. (b) Differential equations for the population dynamics in each phenotype and (c) their analytical solutions. (d) Typical shapes of viable cell growth curve depending on the phenotype rates of changes, *k_P_*, *k_d_*,*s*.

Here, the subscript *m* means a maximum effect for each parameter; e.g., *k_md_* is the maximum effect for *k_d_*, relatively to the normal growth rate, *k*_0_. The growth rate at drug-less condition (*k*_0_) and the intrinsic doubling time (*T_d_*) are related by the relation, *k*_0_ · *T_d_* = ln2. HC_50_ is the concentration corresponding to the half maximal effect for each parameter. These parameter sets determine the growth behavior of each phenotype completely according to the phenotype population dynamics (Figure 1b, c). The growth curve was presented as a fold change (*N_c_*) of viable cells, that is a sum of proliferating cell (*n_P_*) and senescent cell (*n_s_*) at a certain dose *C*. *N*_0_ is the fold change in normal condition, i.e., a drug-less condition (*C*=0).

#### Does-response Curve

For given growth curves for various doses, 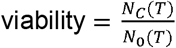 and apparent growth rate 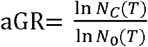 were calculated at a certain time point (*T*), and these calculations generated a dose-response curve. Here, the time difference between the drug injection and the time point at which viability or aGR is calculated, corresponds to the assay duration. Several dose-response curves were generated along the assay duration.

#### Response Summary

For a given dose-response curve of either viability or apparent growth rate, potency (IC_50_ and EC_50_) and efficacy (E_max_) were obtained by nonlinear fitting with a classical Hill function, 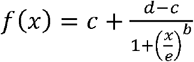. This was performed with the R package ‘drm’. The fits produced well-converged results except in high doses where the growth is too small. The fitting parameter *b* and *d* was 1 in all cases. The summary factors were determined through

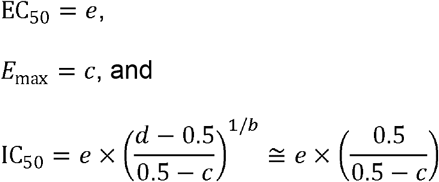

The overall effectiveness was measured as the area under the inhibitory curve (=1-viability), AUC, by using ‘trapz’ function in R package ‘pracma’.

### 2.2 Drug Response Assessment of Public Data

#### Public Data

Drug response data were downloaded from the source data link (https://www.nature.com/articles/s41467-019-10460-1#Sec32) of Fleury’s paper (Fleury et al., 2019). This was the only publicly available dataset to date that included both time-lapse enumerations of viable cells and the end-point measurements of the phenotype fractions. We assessed drug responses of OV1369(R2) and OV1946 cell lines treated with Olaparib through both of the metrics: the conventional metric and the alternative phenotype metric. All data analyses and plots were performed in R.

#### Growth Rates and Viability from Kinetic Measurements

The viable cell counts, divided by the initial number *n*_0_, were applied to a 2-parameter exponential function, *A*(*e^kt^* − 1) + 1. The nonlinear least-squares regression was performed with “nls” function in R. The proliferating-growth-rate, *k*, is given as one of the fitting parameters. For the conventional assessment, apparent growth rate was determined by either nonlinear fitting with a simple exponential function or calculating 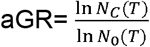 for the assay duration of 3-days and 6-days. 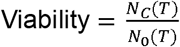 were measured for the same assay duration.

#### Phenotype Fraction

According to the explicit function of each phenotypic growth, the fractions of phenotype increase are given as 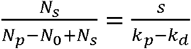 for senescent cells and 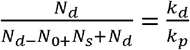 for dead cells. Note that *N_P_* − *N*_0_, instead of *N_P_*, makes the fractions a simple ratio of the parameters. However, the phenotype fractions in the original data are the conventional forms measured by either senescence-associated β-galactosidase assay (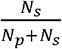 for senescent cells) or flow cytometry(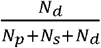 for dead cells). To convert the conventional phenotype fraction to the fraction of phenotype increase, a fold change (FC) of viable cells was used: 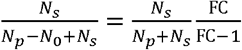 and 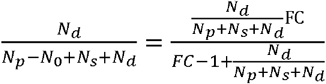 for 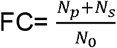 obtained from the kinetic measurements.

#### Phenotype Parameter and Response Classification

*k_P_*, *k_d_*, and *s* were determined by the system of linear equations; proliferating-growth-rate *k* = *k_P_* − *k_d_* − *s*, and the fractions of phenotype increase 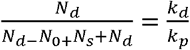, and 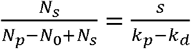 Thereafter, at a certain drug concentration *C*, the variation extent of each parameter relative to *k_P_*(0) ≈ *k*_0_ were calculated. For example, the relative change of *k_d_* is 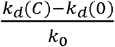. By comparing the relative change of each parameter, a dominant mechanism underlying growth inhibition could be determined.

## 3 RESULTS

### 3.1 Phenotype dynamics model: Each cellular response phenotype contributes differently to the overall growth behavior

Phenotypes related to population growth can be divided into three groups: proliferating cells, dead cells, and cell cycle arrested cells (i.e., senescent cells). Although senescence has never been considered individually in the conventional assessment of drug response, therapy-induced senescence is a widely reported phenomenon (Ewald et al., 2010; Mikuła-Pietrasik et al., 2020; Wang et al., 2020a). The number of cells in each phenotype (*n_P_* proliferating cells, *n_s_* senescent cells, and *n_d_* dead cells) changes through cell division (with rate of change, *k_P_*) or cell death (*k_d_*), or due to cell cycle arrest (*s*) among proliferating cells (Figure 1a). Accordingly, the complete set of differential equations for phenotype population dynamics is given, as shown in Figure 1b, and the explicit analytical solution (Figure 1c) is derived directly without numerical calculation. The death of senescent cells has been disregarded herein, the exception being the coexistence of senolytic drugs or immune cells.

The solution shows several notable points directly. All phenotypes have the same characteristic exponent in their growth function: *k* = *k_P_* − *k_d_* − *s*, that is the growth rate of proliferating cells. We call this just “growth rate (GR)” as distinct from “apparent growth rate (aGR)” that assumes a simple exponential growth of the total viable cells. Viable cell, *n_P_* + *n_s_*, does not follow a simple exponential form. Instead, it follows the function of time, *A*(*e^kt^* − 1) + 1. Depending on *k_P_*, *k_d_*, and *s*, the shape of a growth curve changes from an increasing convex type (*k_P_* − *k_d_* > *s* > 0) to linear (*k_P_* − *k_d_* − *s* ≪ 1) or even concave (*s* > *k_P_* − *k_d_* > 0), and to a decreasing convex type (*k_P_* − *k_d_* < 0) (Figure 1d). However, it can be a simple exponential curve if s~0, that is, for the case of negligible senescence.

### 3.2 Different length of assay and different metrics produce different dose-response curves with the same data

By using the population dynamics of viable cells, we explored how the conventional dose-response curve changes depending on the assay duration and the metric used in producing a dose-response curve. To generate growth curves for certain drug-cell line pairs, we assigned the phenotypic rates of changes for two cases depending on how dominantly the senescence occurs: i) senescence-dominant, or ii) -negligible. The phenotype rates of changes as a function of dose and the growth curves at several doses were presented in Figure 2a. By using these same growth curves, we constructed dose-response curves of viability, aGR, and also GR at different end timepoints of the assay (Figure 2b). If dose-response curves are overlapped together, it means that the used metric for such dose-response curves does not depend on assay duration.

**Figure 2.**
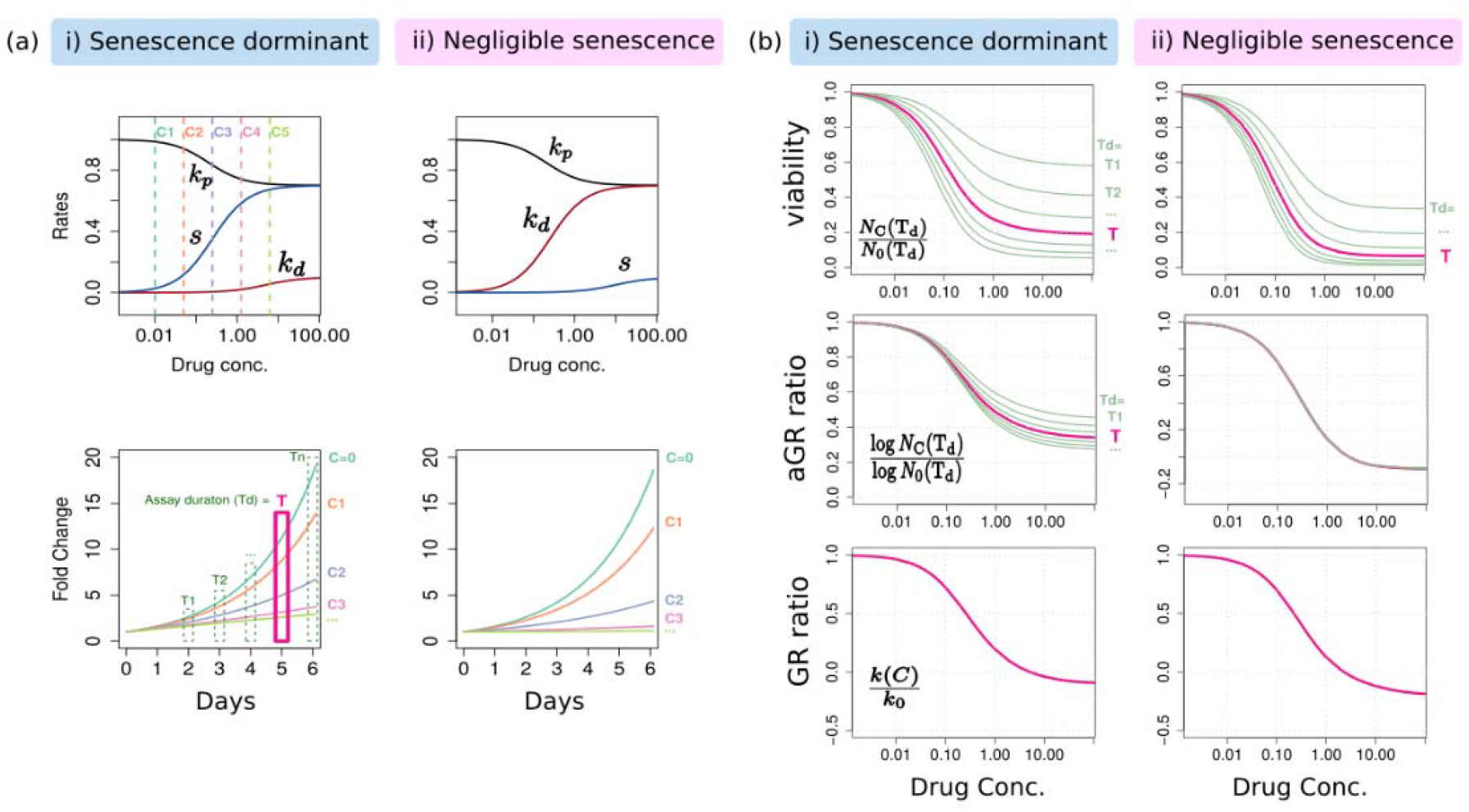
The dose-response curves of two distinct phenotype response: i) senescence-dominant and ii) negligible-senescence. (a) The rates of changes and the corresponding growth curves. The normal growth rate *k*_0_ was assumed as 0.5 (that is, a doubling time =1.4 days). (b) The dose-response curves of viability, aGR ratio and also GR ratio.

Different assay durations produce different dose-response curves of viability; This indicates a significant time-dependency regardless of how significantly senescence occurs (Figure 2b upper). Both potency and efficacy are changed systematically; the longer duration gives the more sensitive response. However, aGR responses is time-independent when senescence is negligible (Figure 2b middle). Otherwise, it depended on time, but less than viability-dose-response did. As a matter of course, GR (that is, *k_P_* − *k_d_* − *s*) was time-independent (Figure 2b lower).

### 3.3 Assay-duration-dependency of drug response causes significant uncertainties in summary factors

Next, we considered what does this time-dependency of the dose-response curve imply in a comparative study of drug response, e.g., a situation of categorizing cell lines to either sensitive or resistant upon a certain drug. The notable point regarding the assay duration dependency is that each cell line is differently influenced by even the same duration of treatment depending on its own doubling time. This can be clearly shown in comparing dose-response curves of two example cell lines that show the same response to a certain drug in terms of cell cycle (Figure 3b): that is, two cell lines having the same GR ratio (of senescence dominant in Figure 2a, i) but different doubling times (either ~1.4 days or ~3.5 days). For the same assay duration, the cell line having the shorter intrinsic doubling time produced the more sensitive response curve (Figure 3b). Therefore, variability in the doubling time of cell lines and a time-dependency of the dose-response make a comparative multi-assay difficult to be controlled consistently in terms of assay length. Instead, a dose-response curve of each cell line has biases relative to each other, that is, there exists intrinsic uncertainty in drug responses.

**Figure 3.**
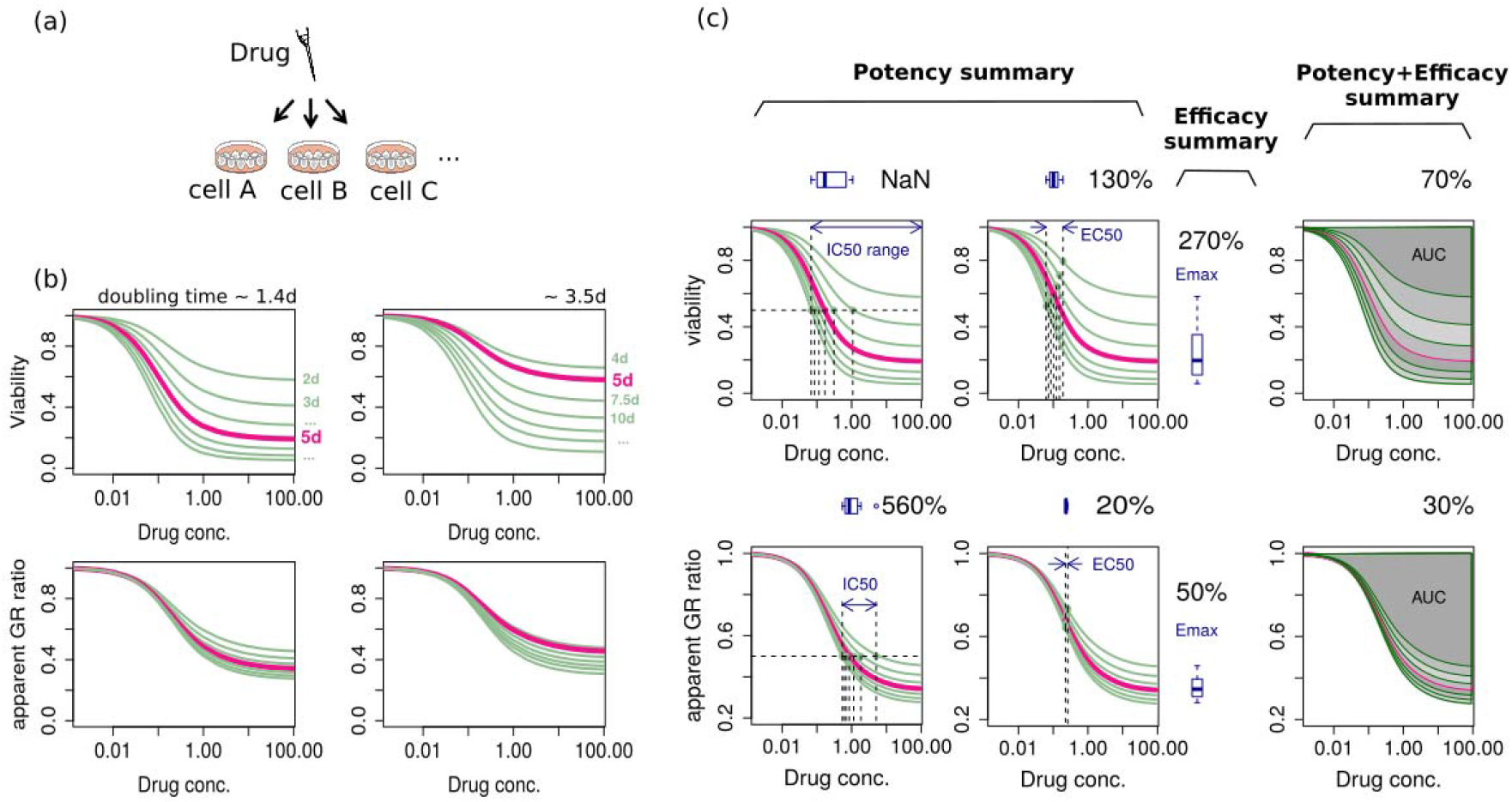
Time-dependency of drug response. (a) A diagrammatic illustration of the response classification assay of various cell lines upon a certain drug. (b) Drug responses of two cell lines that have the same GR ratio but different doubling times. (c) Variation of summary factors (IC_50_, EC_50_, E_max_, AUC) along the assay duration. The box plot, along with the ratio of the maximum deviation to the mean, shows uncertainty of summary factors by time-dependency.

To estimate uncertainty of the summary factors due to time-dependency in a typical experimental condition, we summarized dose-response curves of various assay duration into IC_50_, EC_50_, E_max_, and AUC (Figure 3c). The duration ranged from 2 to 8 days correspond to 1.5~6 times of the original doubling time, 1.4 days, of the cell having *k*_0_=0.5. As a matter of course, viability-based metrics have larger uncertainty than aGR-based one (Figure 3c). Among the summary factors, IC_50_ is largely deviated along the assay duration, even to the infinite in 2-days assay. AUC shows a relatively small deviation in both viability- and aGR dose-response curve.

### 3.4 Effectiveness ranking based on a dose-response curve depends on the assessment method

We considered the second situation of drug screening; assessing therapeutic effectiveness of various drugs for a certain cell line. In this multi-assay, there is no variability of intrinsic doubling time. The question would be either which drugs are effective and which are not, or how to determine the most effective drug to a given cell line? The common way to answer this question is ordering the summary factor. To this end, we investigated therapeutic effectiveness of 7 example drugs with various therapeutic effects in terms of dominant phenotype and potency and efficacy. Summary factors were extracted from each dose-response curve, and displayed in bar plots beside the corresponding response curve (Figure 4). While the lower value of IC_50_, EC_50_ and E_max_ corresponds to the more sensitive response, for AUC the higher value is the more sensitive.

**Figure 4.**
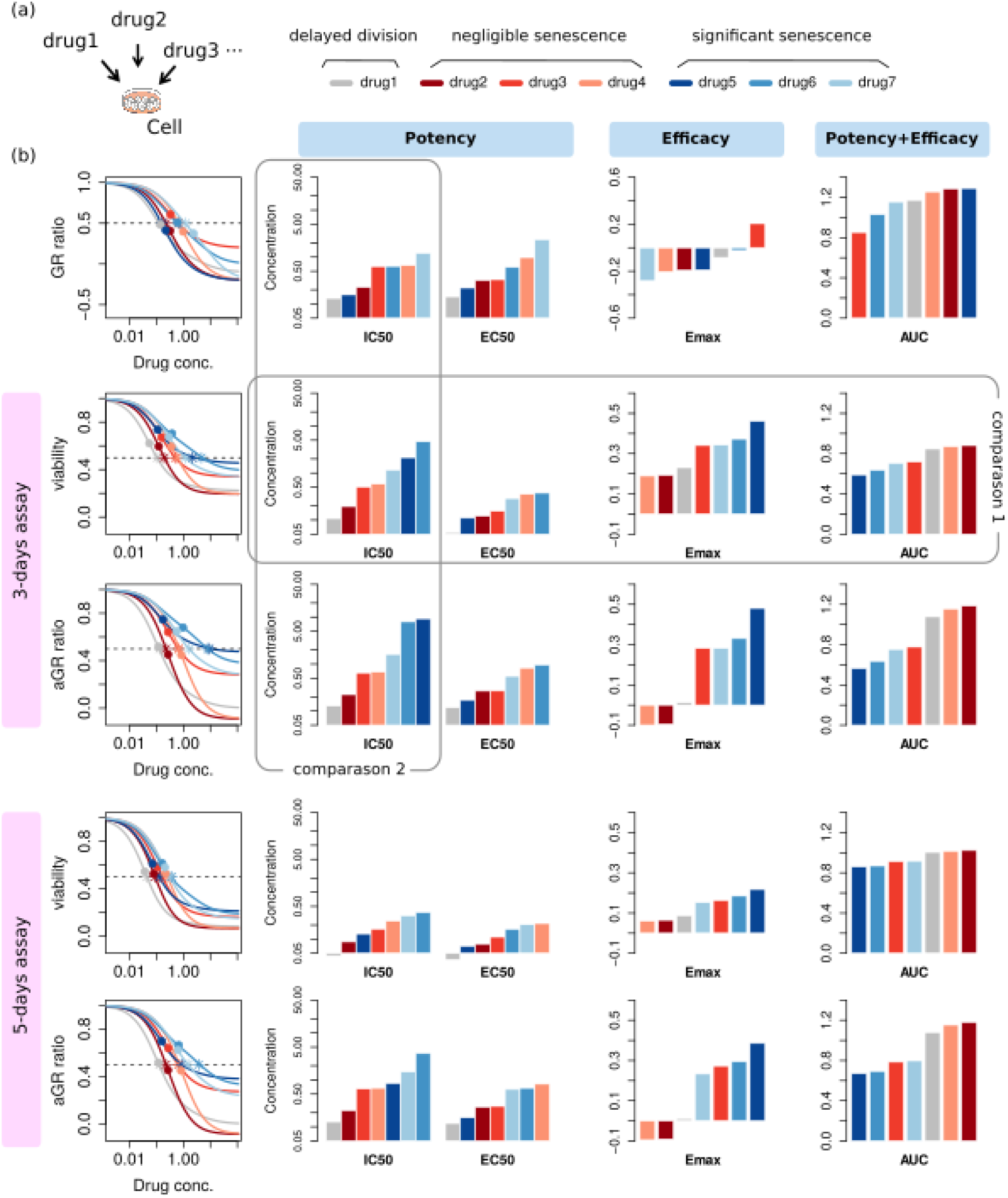
Ambiguity in assessing therapeutic effectiveness by convention metrics. (a) A diagrammatic illustration of the assay. (b) For each dose-response curve, the summary factors, IC_50_, EC_50_, E_max_, and AUC, were extracted. The stars and the circles on dose-response curves correspond to IC_50_ and EC_50_ respectively.

Overall, the assessment result for therapeutic effectiveness shows considerable ambiguity. For example, different summary factors produce different orders for therapeutic effectiveness: According to IC_50_ and EC_50_, the most effective drug is drug 1 (grey), while it is drug 4 (apricot) by E_max_, and drug 2 (dark red) by AUC based on 3-days viability-dose-response curve (Figure 4, comparison 1). Even with the same summary factor, an effectiveness ranking of a drug is different depending on which metric is used for producing a dose-response curve: When summarized into IC_50_, 3-days viability-dose-response curve gives drug 6 (blue) as the least effective one among 7 drugs while aGR- gives the same drug as 6th and GR- gives as 5th (Figure 4, comparison 2). Moreover, even with the same metric and the same summary factor, again a different assay duration produces a different order for therapeutic effectiveness. (Figure 4, 3-days vs. 5-days)

### 3.5 Alternative phenotype metric provides time-independent characteristic quantities of drug response

The above results indicate that the conventional drug response metrics have the significant limitations for comparative analysis of drug response. So, here, we suggest measuring phenotypic rates of change at a certain dose rather than constructing a full-range dose-response curve. Dead cell count to the total increased number of cells 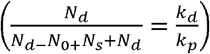, and senescence cell count to total increased number of viable cells 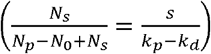, along with GR, provide the system of linear equations that would enable the determination of all three unknown *k_P_*, *k_d_*, *s*.

These findings were confirmed by evaluating real data. We used a publicly available dataset that included both time-lapse enumerations of viable cells and end-point measurements of the phenotype fractions; the ovarian cancer cell lines OV1369(R2) and OV1946 were treated with the PARP inhibitor Olaparib for 6 days (Fleury et al., 2019). At first, dose-response curves of viability and aGR displayed significant differences between 3-days and 6-days assays, indicating their time-dependency (Figure S1 and Supplementary Table 1a, b). IC_50_ of 6-days assays were much lower than those of 3-days assays (1.5–3 times for OV1369(R2) and ~10 times for OV1946).

For the phenotype metric, GR was obtained by nonlinear fitting of the viable cell count. For the growth curves of the lowest 3~4 doses, the fit was converged well. The fraction of phenotype increase was obtained by reformulating the conventional phenotype fraction measured by β-Gal staining or flow cytometry (Figure 5). In the original paper (Fleury et al., 2019), the phenotype fractions were measured just at 4 doses. Therefore, the phenotype parameters could be determined at a single concentration point where both GR and phenotype fractions were determined, along with at the drugless condition (Tables in Figure 5). At that concentration, 2.5μM for OV1369(R2) and 0.01μM for OV1946, Olaparib showed cytostatic effect to each cell line. If we would focus on the response at 1μM, OV1369(R2) is resistant according to whether or not k/k0 is larger than 0.5 at that concentration. In contrast, OV1946 is sensitive because *k*/*k*_0_=0.202/0.534 is smaller than 0.5 even at much lower dose (0.01μM). The dominant mechanism underlying growth inhibition can be classified according to the variation extent of each phenotype rate relative to the normal growth rate. In the response of OV1396(R2) upon Olaparib treatment, senescence is dominant (10.3% for senescence > 0.3% for cell death) in addition to delayed cell division. Regarding OV1946 cells, not only is growth significantly inhibited even at a low dose of Olaparib, but also that this effect results from cell death rather than senescence (26.3% for cell death > 3.8% for senescence)

**Figure 5.**
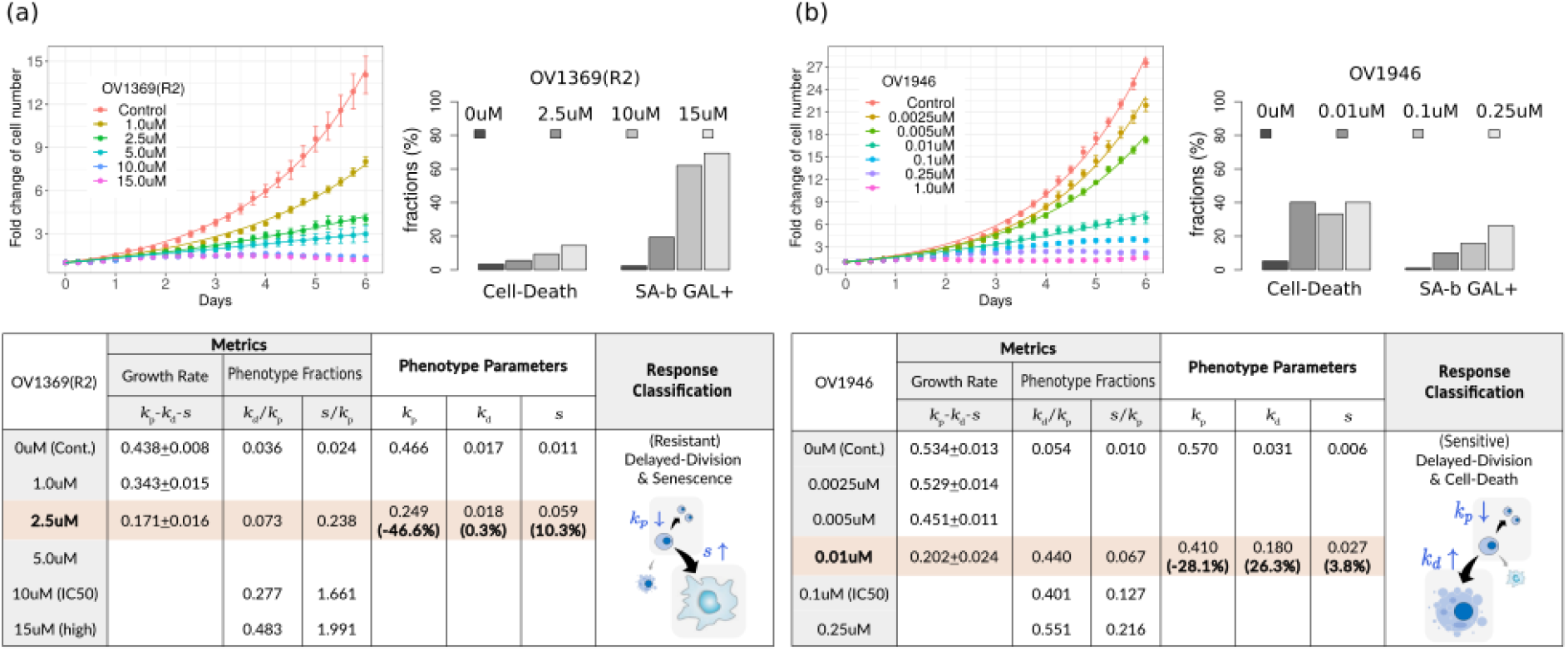
Evaluation of phenotype metric for ovarian cancer cell lines (a) OV1369(R2) and (b) OV1946 treated with Olaparib. Fold change of viable cells and the fraction of dead and senescent cells were re-plotted using the public raw data (upper). Calculation of the phenotype parameters and classification of drug-response were summarized in the table (lower).

## 4 DISCUSSION

We explored two different types of drug response metric under the consideration of phenotypic population dynamics: the conventional one based on a dose-response curve and the alternative phenotype metric evaluated at a single dose.

The drug response has been typically represented by summary factors extracted from a dose-response curve. The most widely used metric is viability. However, the difference in cell counts between drug-treated and -untreated condition increases with time; that is, a longer treatment results in lower cell viability. This tendency becomes more prominent with an increase in dosage; hence, the longer the assay duration, the more sensitive the dose-response curve. Moreover, the assay duration is a relative factor for the intrinsic doubling time of each cell line, which means that a viability-based metric has a serious limitation for comparing the drug responses of various cell lines. In this regard, the growth rate could be a reasonable alternative to viability (Hafner et al., 2016; Harris et al., 2016).

However, if the growth rate is derived on the assumption of a simple exponential growth even under drug treatment, the dose-response curve might still depend on time. A simple exponential growth results from the premise that the rate of cell number changes is proportional to the number of cells in each given timepoint. The proportional factor is growth rate. If some of the viable cells are undergoing cell cycle arrest–which does not contribute to changes in the cell count–viable cells do not follow simple exponential growth. The number of proliferating cells itself might be an exponential function; however, the total number of viable cells (including both proliferating cells and cell cycle-arrested cells) is not. This indicates the need for considering the actual cellular response phenotypes when evaluating drug responses.

In this regard, we considered how each cellular response phenotype contributes to the population dynamics. Indeed, the viable cell count was not a simple exponential function but *A*(*e^kt^* − 1) + 1. It indicates time-dependency of the aGR is intrinsic because of the senescent cell subpopulations regardless of other possible confounders mentioned in the literature (Niepel et al., 2019). On the contrary, GR (*k*= *k_P_* − *k_d_* − *s*) is time-independent, and it is the only characteristic exponent in the population dynamics. Unless specifically enumerating the proliferating cells, a proper fitting model, not a simple exponential function, should be applied to the number of either viable cells 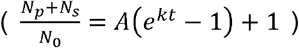 or dead cells 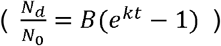 to determine GR. In therapeutically effective dose where the growth is too small, measuring dead cell counts would be better for the convergence of nonlinear fitting. Recent advancements in time-lapse imaging in live cell microscopy (Gelles et al., 2019) potentially provide a useful platform for measuring the cellular kinetics to determine the growth rate.

Through properly figured population dynamics, the conventional drug response metrics showed several limitations. In the case of a response classification of cell lines, the summary factors have a significant uncertainty because of the assay-duration dependency of the dose-response curve and doubling-time variability of cell lines. This is an additional uncertainty to other experimental variabilities such as seeding numbers, effects from either a direct cell counting or CellTiter-Glo assay, and edge effects in a well plate (Hafner et al., 2016; Niepel et al., 2019). Obviously, summary factors from an aGR-based dose-response curve have a smaller uncertainty than those from a viability-based curve. Among the widely used summary factors, EC_50_ has the smallest deviation but it does not represent an overall effectiveness, only potency. Efficacy is captured separately as E_max_. Instead, AUC captures an overall effectiveness as well as showing a relatively small deviation. The most common factor, IC_50_, is the worst in terms of uncertainty as shown in a statistical framework study that uses uncertainty estimations to improve biomarker discovery with assessment of cell line drug response (Wang et al., 2020b).

Even in the case of drug screening for a single cell line wherein there is no doubling time variability, the dose-response curve-based assessment has intrinsic ambiguity in that the order of the therapeutic effectiveness changes depending on the metric (viability or aGR or even GR) for a dose-response curve and the summary factor, and the length of the assay in case of the conventional metric. The ambiguity of therapeutic effectiveness assessment using a dose-response curve is represented even in the GR-dose-response curve, that is, time-independent, characteristic growth rate. This is intrinsic in that summarizing a dose-response curve is done customarily by fitting with the empirical sigmoidal curve. Without a theoretical basis, the conventional summary factors are an apparent quantity, as a simple exponential fitting gives just an apparent growth rate for viable cell growth, not the characteristic quantity. Furthermore, summarizing to apparent quantities can increase ambiguity more than raw uncertainties in experimental variabilities by uncertainty propagation (Wang et al., 2020b). This might be reasons for not only dimness and obscurity within a pharmacogenomic study but also inconsistency or a modest correlation between large pharmacogenomic studies like CCLE (Barretina et al., 2012), GDSC (Yang et al., 2013), and CTRP (Seashore-Ludlow et al., 2015). This finding is in line with a previous report regarding systematic variations of the summary factors in large scale drug response data (Fallahi-Sichani et al., 2013). In that study, different summary factors captured distinct information and the most informative factor varied with the drug. Therefore, this study concluded that factors other than potency should be considered in the comparative analysis of drug response, particularly at clinically relevant concentrations. However, while the general cellular growth as a function of time is a derivative form of exponential function with a characteristic exponent, how about this characteristic quantity itself as a function of dose? Classical pharmacology is not ready to explain a dose-response with theoretical basis. Unlike in case of growth curves, it is difficult even to expect dose-response curves have a characteristic function or quantity in general.

In this regard, we suggest an alternative evaluation of drug response that is not based on a dose-response curve but focuses on the response at a single dose as it is. However, GR alone is not enough for an assessment of drug response, because there are various combinations of k_P_, k_d_,s for the same GR and we want a response variable and classification to have phenotypic relevance. We need to measure all phenotype rates, that contain all the information of the drug response at a given dose and are directly connected to the mechanism underlying growth inhibition.

This phenotype metric provides clear and phenotype-relevant pharmacogenomic variables. The therapeutic effect is categorized either as cytotoxic or cytostatic, and the dominant mechanism of action either as delayed division, senescence, or death. Only in the case of *k*_P_ − *k_d_* < 0, the effect is cytotoxic (that is, actual decrease of viable cells); otherwise, it is cytostatic (inhibited growth). Even if *k_P_* − *k_d_* − *s* is negative, in the case of (*s* > *k_P_* − *k_d_* > 0), the viable cell number increases and they all finally become senescent cells with a certain saturation level 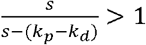. Therefore, the detailed classification is either inhibited growing (*k_P_* − *k_d_* > *s* > 0), expanded saturation (*s* > *k_P_* − *k_d_* > 0), or shrunk saturation (*k_P_* − *k_d_* < 0). Note that, unless s ≈ 0, a complete shrinkage does not happen, because the fold change of viable cells goes to a nonzero level 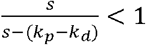. To classify the mechanism underlying growth inhibition, 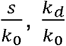 and 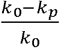 can be compared. The classical categorization into either sensitive or resistant still does not have a theoretical standard, but a *de facto* classification might be applicable in line with the definition of IC_50_; that is, by whether or not the GR ratio (*k*/*k*_0_) is larger than 0.5 at a clinically relevant dose. Evaluation flows for the conventional metrics and the alternative phenotype metric were summarized in Figure 6.

**Figure 6.**
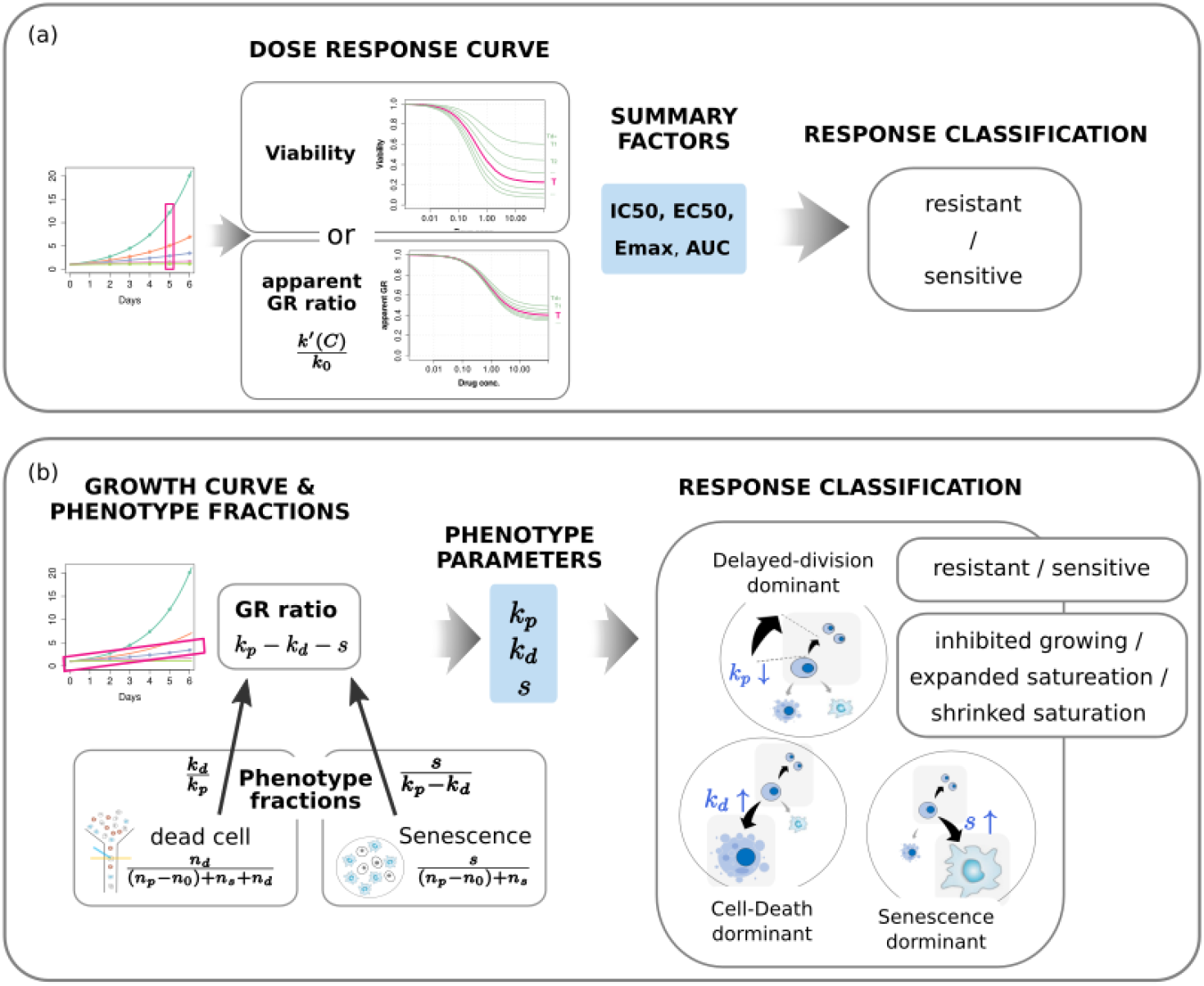
Evaluation of drug response based on (a) the conventional dose-response curve and (b) the phenotype population dynamics at a single dose. Three types of response classification are possible with the phenotype metric.

We believe the ability of the phenotype metric to provide the characteristic quantities of drug response and the mechanism underlying growth inhibition would markedly improve pharmacogenomic analysis. This improvement of the pharmacogenomic variables in preclinical pharmacology will provide better translation into clinical studies and useful information for treatment decision-making. The absence of need for a very wide range assay is an additional advantage of this phenotype metric. A single-concentration assay at a clinically relevant range in addition to no-drug assay is enough. Of course, evaluations in a wider range of concentrations would give a separate dose-response curve for each of *k_P_*, *k_d_*,*s*, that would provide rich information regarding therapeutic mechanisms of action.

Particularly, the significance of senescence in drug response is notable. Deviation of growth behavior from a simple exponential function and no complete shrinkage of viable cells are definitely the consequence of senescence. Furthermore, senescence causes systematic variation in conventional metrics; for the same GR dose-response, the more significant the senescence, the less sensitive the dose-response curve of viability and aGR. If an assay duration increases, then the tendency for a decreased sensitivity along senescence is mitigated. This is because senescence reduces the proliferating growth rate, but it does not reduce viability.

Even though the conventional dose-response curve has some limitations for providing precise pharmacogenomic variables, it is still useful for a qualitative visualization of overall drug responses of a single cell-line upon a certain molecular feature. For example, ML239 cytotoxicity in NCIH661 LCLC cell line along the knockdown efficiency of *FADS2* were well visualized in the viability dose-response curve (Rees et al., 2016). Therefore, we conclude it is appropriate to use the phenotype metric for therapeutic effectiveness assessment related to pharmacogenomic association studies and the dose-response curve for qualitative visualization of overall drug response.

## ACKNOWLEDGMENTS

This research was supported by Basic Science Research Program through the National Research Foundation of Korea (NRF) funded by the Ministry of Education (No. NRF-2019R1A6A1A03032888) and by a grant (HI16C1559) from the Korea Health Technology R&D Project through the Korea Health Industry Development Institute (KHIDI), funded by the Ministry of Health & Welfare, Republic of Korea.

## AUTHOR CONTRIBUTIONS

S. K. and S. H. conceived the idea and wrote the manuscript. S. K. built and evaluated the mathematical models and analyzed the public data. S. H. supervised the study.

## DATA AVAILABILITY

Data sharing is not applicable to this article for no new data was created in this study. The data that support the findings of this study are available in the source data file of Fleury’s work at (https://www.nature.com/articles/s41467-019-10460-1#Sec32).

## ETHICS APPROVAL

Not applicable-no new data generated.

## PERMISSION TO REPRODUCE MATERIAL FROM OTHER SOURCES

Not applicable-no reproduction.

